# Exploring Holography in Neuro-Vascular Dynamics

**DOI:** 10.1101/2025.05.19.654699

**Authors:** Christian Kerskens

## Abstract

The holonomic brain theory—originally formulated to account for the need of non-local memory encoding in cognitive systems—could gain new theoretical traction when integrated with holographic principles from physics, most notably the AdS/CFT correspondence. Recent findings in neuroscience suggest that conformal field theories (CFTs), emerging at critical points across spatiotemporal scales in neural dynamics, are essential for brain function. Concurrently, black-brane geometries, long studied in gravitational physics, can find unexpected analogues in the interplay of active matter dynamics and the brain’s neuroanatomical organization. Motivated by these parallels, we posit a generalized holographic framework and interrogate its validity through the fluid/gravity duality—a correspondence linking hydrodynamic equations to gravitational spacetime metrics. In this work, we explore the holographic principles at the Navier-Stokes regime, demonstrating that holography can model key neurophysiological mechanisms: cerebral autoregulation (the brain’s hemodynamic self-stabilization) and neurovascular coupling (the dynamic neuron-bloodflow interplay). This work bridges holography, active matter physics, and neuroscience, proposing a unified framework to decode the brain’s multiscale organization, its resilience to perturbations, and its computational capabilities. By grounding neurovascular physiology in gravitational duals, we open pathways to reinterpret brain function through the lens of emergent spacetime geometry.

## 1. Introduction

Conformal field theories (CFTs) arise at the critical point of condensed-matter systems, where scale and conformal invariance emerge (for example, the 2D Ising model [1,2] for two-dimensional membranes or thin films). In the brain, cell membranes are believed to function close to criticality in their lipid and protein organization, potentially allowing for their extraordinary adaptability [3]. Lipid bilayers may phase separately into coexisting liquid-ordered and liquid-disordered domains, and experiments carried out on giant plasma membrane vesicles from mammalian cells show that these vesicles often are situated very near to the critical point of a two-dimensional Ising model [4,5]. Near this transition, lipid composition fluctuations, predominantly affecting cholesterol content, are scale invariant, optimizing membrane flexibility and promoting dynamic protein clustering and signaling. Simultaneously, ion channels that sit within these fluctuating lipid environments might harness criticality to enhance their sensitivity. Cooperative interactions between specific lipids and channel proteins (for example, voltage-gated potassium channels) exhibit signatures of critical dynamics, allowing membranes to modulate gating sensitivity in response to small perturbations [6].

Likewise, a significant number of membrane receptors are thought to use critical-like transitions within their conformational landscapes to amplify signals; even small perturbations in ligand binding or lipid packing can yield large changes in activity in these systems [7]. Recent theoretical and computational studies have focused on determining the scaling dimensions and operator content governing membrane phase separation, as well as predicting critical Casimir forces between membrane inclusions such as proteins [3,8]. These long-ranged, fluctuation-mediated forces can drive weak aggregation or repulsion of membrane proteins, with potential implications for signaling platforms and cytoskeletal attachments. Indeed, critical phenomena are observed in the brain far beyond microscopic scales such as cell membranes — they appear to underlie organization at every level of analysis, from individual synapses to whole-brain networks.

Spontaneous activity observed in neocortical tissue [9,10], known as “neuronal avalanches”, is characterized by scale-invariant power-law distributions in both the size and duration of bursts, a signature which has been argued to optimize information transport and expand the range of dynamic network behaviour. At the level of synapses, long-term potentiation and depression are in mutual balance, spontaneously pushing synaptic strengths to a critical state, which provides maximal memory capacity yet maintaining stability of the network [11].

As we proceed to larger circuits, the brain continuously shifts between synchronized and desynchronized states across multiple frequency bands. For example, beta rhythms synchronize over longer distances than gamma rhythms, indicating distinct roles in long-range versus local communication [12]. Furthermore, evidence of critical dynamics spans a broad frequency spectrum—from slow fluctuations around 0.05 Hz up through fast oscillations near 125 Hz—highlighting criticality as a unifying principle of brain function [13,14]. This balanced state allows rapid and quick coordination of activity to external requirements.

Anatomically and functionally, the networks of the brain have fractal and scale-invariant structures, which mediate information flow effectively across spatial scales. Underpinning these dynamics are homeostatic mechanisms—from astrocyte-mediated regulation to neuromodulatory systems such as dopamine—that constantly adjust neuronal excitability so that neither runaway excitation nor silence prevail [15]. Taken together, these multiscale observations suggest that critical dynamics are not mere curiosities, but they play an essential role in the brain and are the source of the impressive adaptability, efficiency, and resilience of neural systems. Further, the brain may exploit the universal physical principles, which CFTs exponentiate, to optimize their scaling and structure and their functional responsiveness.

There are also indications that those CFTs and the brain as a whole are holographic. In neuroscience, the holonomic brain theory, proposed by Karl Pribram, suggests that memory is encoded in wave interference patterns within dendritic webs, analogous to holographic storage [16,17]. This model explains non-local memory storage and fast associative recall. At tissue level, observations of quantum entanglement have been reported in the awake brain [18], which suggests exotic physics, like CFTs, at play. At the Navier-Stokes level, features specific to CFTs at cell level are carried on to fluid dynamics as demonstrated in resting-state fMRI, which shows 1/f-type temporal correlations, another hallmark of criticality that facilitates long-range temporal integration [19].

Here, we want to explore this further. The dynamic system of the brain can potentially create Euler-like flow pattern (using active matter to reduce the viscosity [20,21]) in some regimes allowing the derivation of analogue black-brane metrics [22,23], which can be tuned into AdS-Schwarzschild, Lifschitz, and Schrödinger metrics depending on experimental conditions. The question whether a correspondence exist between the brain’s CFTs and the potential black-branes, can be investigated using the fluid/gravity duals [24]. In a holography, the fluid/gravity dual can be seen as the bridge between the boundary’s CFT and the black-brane. Assuming the existence of a holographic framework, a valid fluid/gravity dual is evidential for the presence of a holographic correspondence.

We will compare the three bulk models AdS-Schwarzschild, Lifschitz, and Schrödinger, which can have a fluid/gravity dual, to find out which is most suitable to describe physiology in the fluid/gravity dual. We will first investigate what the likelihood of a strongly-coupled Navier-Stokes system in the living brain is, which is a necessary condition for the investigatd fluid/gravity dual. Then, using the cerebral auto-regulation and the neuro-vascular response, we will narrow down which holography could, if at all, describe a holographic brain.

## 2. Results

### Cerebral autoregulation

is the brain’s ability to preserve cerebral blood flow at a constant level despite changes in blood pressure [25,26]. The mechanisms underlying cerebral autoregulation are complex and not fully understood, with ongoing debate about the role of sympathetic nervous activity [25]. Within a mean arterial pressure range of 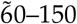 mmHg, flow velocity is relatively stable. Within a mean arterial pressure (MAP) range of 60–150 mmHg, flow velocity is relatively stable. Studies such as those by Aaslid et al. [27] and Paulson et al. [28] demonstrate this auto-regulatory capacity. If we begin with describing cerebral blood flow dynamics by the incompressible Navier-Stokes equation (with constant density *ρ*):

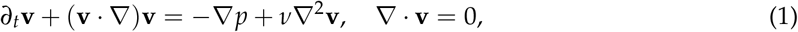

where *ν* = *η*/*ρ* is the kinematic viscosity, we find that under the *z* = 2 dilation

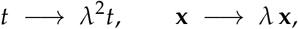

the hydrodynamic fields change according to

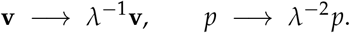

Thus, independent of the state of the incompressible Navier–Stokes equation, every term scales with the same overall order, which means that the equation

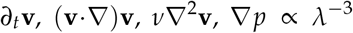

remains form-invariant under the global rescaling. However, the scaling of the pressure gradient, which scales with Δ*p* → *λ*^−3^Δ*p* and the velocity which scales with **v** → *λ*^−1^**v**, indicate that dynamics described by the incompressible Navier-Stokes equations are incompatible with the auto-regulation. By contrast, Navier-Stokes equations show a different behaviour in holographic correspondences.

The fluid/gravity duality itself exhibits that a solution of the Einstein’s equations near an asymptotically AdS black brane leads in the derivative (long-wavelength) expansion to relativistic Navier–Stokes equations on the conformal boundary [24,29]. One can also obtain non-relativistic hydrodynamics via a similar strategy by tuning the structure in the bulk: Lifshitz spacetimes with dynamical exponent *z* realize anisotropic scale invariance, and for *z* = 2 their boundary theory generates the diffusive scaling symmetries of the incompressible Navier–Stokes equation [30]; Schrödinger spacetimes generalize this construction by including a null direction that geometrizes Galilean boosts and a conserved particle number current. At leading order one again recovers a *z* = 2 non-relativistic fluid [31,32].

**Table 1.**
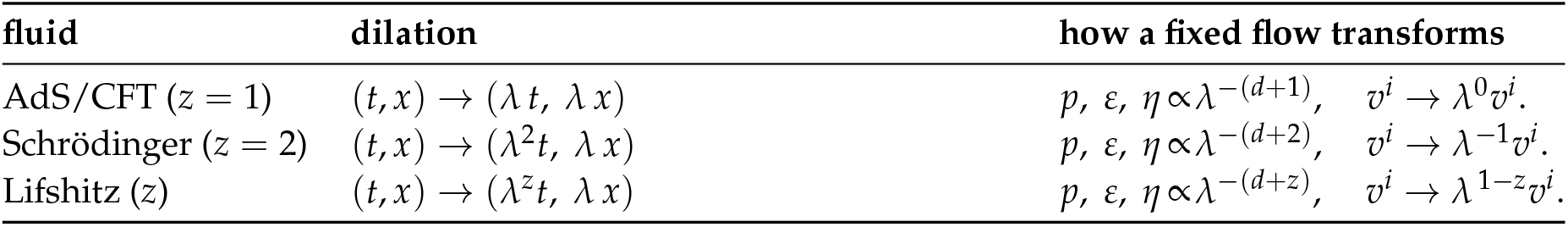
Visualising the scaling test under a global dilation.

| fluid | dilation | how a fixed flow transforms |
| --- | --- | --- |
| AdS/CFT ($z = 1$) | $(t, x) \rightarrow (\lambda t, \lambda x)$ | $p, \varepsilon, \eta \propto \lambda^{-(d+1)}, \quad v^i \rightarrow \lambda^0 v^i.$ |
| Schrödinger ($z = 2$) | $(t, x) \rightarrow (\lambda^2 t, \lambda x)$ | $p, \varepsilon, \eta \propto \lambda^{-(d+2)}, \quad v^i \rightarrow \lambda^{-1} v^i.$ |
| Lifshitz ($z$) | $(t, x) \rightarrow (\lambda^z t, \lambda x)$ | $p, \varepsilon, \eta \propto \lambda^{-(d+z)}, \quad v^i \rightarrow \lambda^{1-z} v^i.$ |

A single global dilation highlights how the three canonical holographic fluids differ.

a. For Schwarzschild–AdS (*z* = 1) the Weyl symmetry of the boundary CFT rescales intensive quantities but leaves the velocity profile intact.
b. For Schrödinger (*z* = 2) fluids a dilation acts as *t* → *λ*^2^*t, x*^*i*^ → *λx*^*i*^, *v*^*i*^ → *λ*^−1^*v*^*i*^; Galilean boosts cannot remove this *λ*^−1^ factor, so the velocity profile itself rescales [33].
c. For Lifshitz fluids the same scaling gives *v*^*i*^ → *λ*^1−*z*^*v*^*i*^; only at *z* = 1 (or with extra symmetries) does the flow remain unchanged [33].

In all three Einstein-gravity cases *η*/*s* = 1/4*π*. Bulk viscosity is zero for relativistic CFTs and Schrödinger fluids, and also for the minimal Lifshitz branes often considered, though it can appear once scaling is broken or higher-derivative terms are added [34].

For the auto-regulation, we can identify the AdS-Schwarzschild as a potential candidate that would literally leave velocity unchanged if the pressure gradient changes. The non-relatistic metrics don’t fulfil this condition. However, it is unlikely, that the local velocity profiles doesn’t change. In medicine, flow velocity are related to a mass flow, mostly described by 1. Fick’ Law [35], which probably means that the velocity invariance doesn’t hold in the brain. For the auto-regulation, we are looking instead for a mass flow which is independent of the pressure gradient in the Navier-Stokes regime. This is exactly the situation in Schrödinger-holography. A Schrödinger-holographic setup includes a null circle *ξ* in the bulk ansatz, which gives the conserved mass (particle-number) current

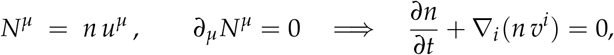

where the boundary particle-number current *N*^*µ*^ is sourced by the bulk null-circle isometry *∂*_*ξ*_ in the Schrödinger geometry, and *n* is the mass density conjugate to the chemical potential. This identification follows directly from the Kaluza–Klein reduction along the null circle *ξ* in the bulk Schrödinger metric [36].

In summary, we found two holographic set ups, which leave “flow” invariant. The Schrödinger holography is probably can provide the most realistic interpretation. However, we will keep the AdS case in our further consideration.

### Neurovascular coupling

refers to the complex communication between neurons, astrocytes, and cerebral vessels that adjusts blood flow to meet the energy needs of activated neurons [37]. While neuronal activation typically precedes vascular changes, the exact timing and mechanisms vary. Fast neuronal Ca2+ transients occur within milliseconds of stimulation, followed by hemodynamic responses seconds later [38]. Astrocytes may play a modulatory role, with their Ca2+ responses occurring between neuronal and vascular changes [38]. The vascular response can include both early and late components, potentially involving different mechanisms [39].

In a holographic picture, like the AdS/CFT the CFT “lives” on the conformal boundary at *r* → ∞, while the Navier-Stokes (NS) fluid appears very close to the black-brane horizon at *r* = *r*_*h* through the fluid/gravity or membrane paradigm. If we translate this situation to the brain, then the cell membranes or similar are at criticality portraying the CFT and the vascular response portraying the NS regime. The huge gravitational red-shift between those two radial locations means that the boundary time *t* and the proper (or EF) time *τ* experienced by the horizon fluid differ by an exponential factor. A signal that is injected at the boundary therefore needs a logarithmically long interval in *t* before it can influence the horizon degrees of freedom that obey the NS equations. This is indeed what we observe in the brain too. The question is if the observable times in a holography system can be as long as in the brain which depending on the species can extend to several seconds.

Consider a planar AdS_*d*+1_ Schwarzschild (black–brane) metric in Poincaré coordinates with the radial coordinate *r*:

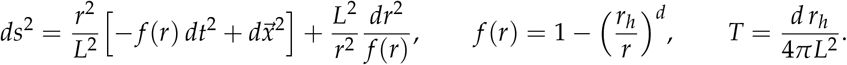

A radial null ray satisfies *dt*/*dr* = *L*^2^/ (*r*^2^ *f* (*r*)). The coordinate time elapsed between the boundary *r* = ∞ and a cut-off surface *r* = *r*_*h*_ + *ε* (the “stretched horizon”) is therefore

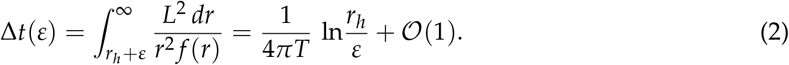

The integral converges in the UV; the large piece is the log IR divergence from the neighbourhood of the horizon *f* (*r*) ~ *d ε*/*r*_*h*_. Equation (2) is universal for any *d* because only the simple zero of *f* (*r*) matters. A change imposed at the boundary takes, in boundary time,

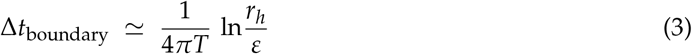

to be felt on a surface a proper distance *ε* outside the horizon. At the exact horizon *ε* → 0 the delay is infinite—the familiar infinite red-shift. The incompressible NS equations emerge if we choose the cut-off to sit parametrically close to the horizon:

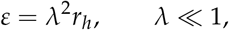

and simultaneously zoom the boundary coordinates as *x*^*i*^ → *x*^*i*^ /*λ, t* → *t*/*λ*^2^. This is exactly the near-horizon/long-wavelength scaling of Bredberg-Keeler-Lysov-Strominger and many later works. Using *ε* = *r*_*h*_*λ*^2^ in (3) gives

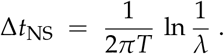

For any fixed but small *λ* (say *λ* = 10^−2^) the fluid variables respond after Δ*t*_NS_ ~(1/2*πT*) × 4.6. For a fluid model, this delay is in the millisecond range to small to explain the vascular response.

Moving on to Schrödinger Holography (Non-relativistic, *z* = 2). One obtains the Schrödinger space-time with anisotropic scaling

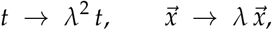

and a null “internal” circle *ξ* by a null Melvin (TsT) twist of AdS_*d*+3_ [36]. In *d* + 3 bulk dimensions (with *d* nonrelativistic spatial boundary directions) the black-brane metric reads

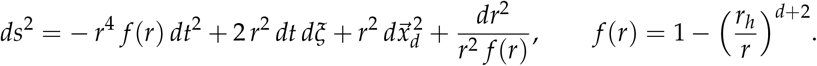

The Hawking temperature is

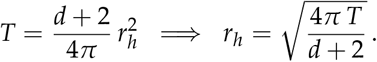

Rather than following null geodesics, the relevant time scale is set by the lowest *non-hydrodynamic* quasinormal mode (QNM) in this geometry. Consider a fluctuation

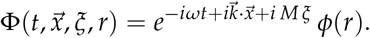

In the dimensionless variables

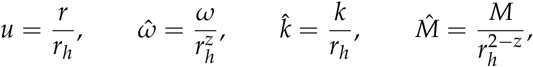

the radial ODE for *ϕ*(*u*) depends only on 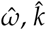 and 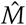. Imposing ingoing boundary conditions at the horizon (*u* = 1) and normalizability at the boundary (*u* → 0) yields a discrete spectrum 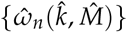. The first non-hydrodynamic mode 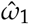 then gives

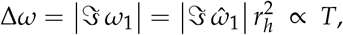

with 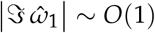. Hence the *thermal delay* is

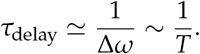

Signals in Schrödinger holography generically carry momentum *M* along the null circle *ξ*. Dropping 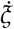 entirely (i.e. setting *dξ* = 0) misrepresents both signal propagation and the QNM spectrum. A proper treatment keeps *M* ≠ 0 in the fluctuation ansatz or geodesic equation—but in either case the hydrodynamic “turn-on” is controlled by the QNM gap, not by a diverging null-geodesic integral. However, Schrödinger holography allows one to engineer a *tunable* signal delay by compactifying the null direction [40]:

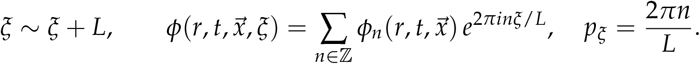

In the uncapped geometry this leads to an infinite-image two-point function [41]:

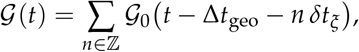

where Δ*t*_geo_ is the single-winding geodesic delay and

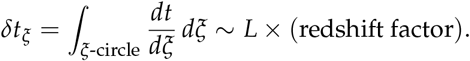

By (i) setting the *n* = 0 source to zero—so no neutral mode is excited—and (ii) smoothly “capping off” the *ξ*-circle at *r* = *r*_0_ (imposing *g*_*ξξ*_ (*r*_0_) = 0 and fixing *L* by regularity) [42], one lifts all |*n*| ≥ 2 modes to large mass, leaving only the |*n*| = 1 images. The net two-point function collapses to

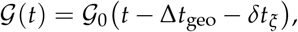

giving a single, *tunable* delay *δt*_*ξ*_ ~ 1/*r*_0_.

Through the fluid–gravity correspondence [43], this translates directly into a strictly silent period

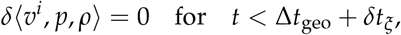

after which the first Navier–Stokes response appears sharply at

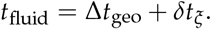

This “dial-a-delay” effect—controlled geometrically operates independently of microscopic transport coefficients (*ν, ζ*) or initial data. Demanding smoothness of the (*r, ξ*) cigar at *r* = *r*_0_ gives

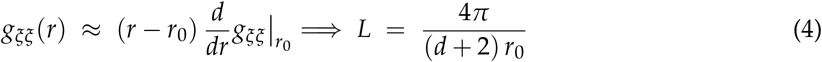

resulting in the fixed-geometry delay

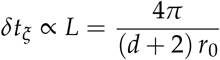

is set entirely by *r*_0_ (and the background redshift), and is blind to *ν, ζ*, or your choice of boundary source profile. The impulsive boundary source induces a **sudden but finite** acceleration in the fluid, governed by the Navier-Stokes equations. While the velocity field itself develops a discontinuity at *t*_fluid_, its time derivative becomes singular but regularized by viscosity:

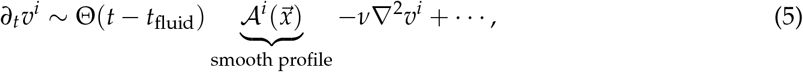

where Θ(*t* − *t*_fluid_) is the Heaviside step function enforcing causality, 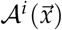 encodes spatial structure from the UV source, and (*ν*∇^2^*v*^*i*^) is the viscous damping smoothing the sharp onset over a timescale *τ*_damp_ ~ *ν*^−1^.

Though geometrically delayed by *L*, the magnitude of the fluid response still depends on transport coefficients (*ν, ζ*), while the timing (*t*_fluid_) remains fully geometric. This resolves the apparent tension: viscosity governs how the fluid moves, but when it moves is fixed by *L* [31,32].

Equation (5) shows the UV-IR interplay: Geometric delays imprint directly on hydrodynamic acceleration, not just propagators. The *L*-dependence in (4) demonstrates how bulk topology regulates non-equilibrium fluidization timescales.

In a water-wave analogue of Schrödinger holography, the bulk null circle *ξ* is mimicked by either (i) a closed-loop flume of circumference *C* or (ii) standing-wave modes in a finite tank. The discrete, KK-like spectrum *k*_*n*_ = 2*πn*/*C* emerges from the periodic boundary conditions [22,23,44]. The effective Hawking temperature in such systems is *T*_H_ ~ 10^−7^–10^−6^ K, giving phonon correlation times of order

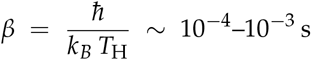

[22,23]. A perturbation propagates around the flume with

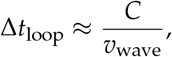

so for *C* = 10 m and *v*_wave_ = 0.5 m/s one finds

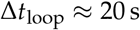

[45]. In a linear tank of length *L*, reflections give

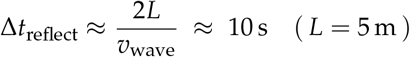

[44].

These two mechanisms—analogue Hawking-temperature decorrelation and geometric loop delays—together capture the “dial-a-delay” of Schrödinger holography in a tabletop water-wave system, entirely independent of fluid viscosity or the initial wave-packet shape. This shows that a translation to a cerebral model could also, in principle, produce delays in the range of circulation times, which makes it then physiologically relevant.

## 3. Discussion

A controlled holographic expansion requires a large-*N* limit, where *N* indexes either the rank of an internal gauge group [46] or the number of components in an vector model [47], which, at first sight, is not achievable in biology.

Nevertheless, holography, particularly through the framework of the AdS/CFT correspondence, has become a valuable tool in studying scaling processes in condensed matter systems [48], even without proof of the framework’s existence.

We applied the same approach to the brain. Rather than proving, we have collected further evidence that the brain uses a holographic framework. We found not only evidence that the fluid/gravity dual is valid, we could also show that two fundamental principles in brain physiology, can be explained with a Schrödinger holography.

Finally, we want to mention that the fundamental principle behind memory and consciousness are still unknown. Holographic principles, as proposed here, can open up new avenues and build on work around the information paradox.

In summary, we have seen that hydrodynamics in the brain may capture how the boundary stress tensor evolves (fluid/gravity) [24,43]. Previously, we found quantum entanglement in the brain, which tells us how changes in boundary entanglement reshape the bulk metric [18,49,50].

Demanding consistency of both (a) the stress-tensor dynamics and (b) the entanglement first law deriving from the same bulk metric would force the nonlinear Einstein equations themselves [51]. Together, they imply that gravity—Einstein’s equations, horizons, area laws—emerges as the universal hydrodynamics of quantum entanglement [52]. The findings in the brain would elevate the holographic principle from a string-theory conjecture to a universal, emergent correspondence [53].

## Acknowledgments

This project has been supported by Trinity College Institute of Neuroscience

## Disclaimer/Publisher’s Note

The statements, opinions and data contained in all publications are solely those of the individual author(s) and contributor(s) and not of MDPI and/or the editor(s). MDPI and/or the editor(s) disclaim responsibility for any injury to people or property resulting from any ideas, methods, instructions or products referred to in the content.

